# High content analysis of granuloma histology and neutrophilic inflammation in adult zebrafish infected with *Mycobacterium marinum*

**DOI:** 10.1101/762740

**Authors:** Tina Cheng, Julia Y Kam, Matt D Johansen, Stefan H Oehlers

## Abstract

Infection of zebrafish with natural pathogen *Mycobacterium marinum* is a useful surrogate for studying the human granulomatous inflammatory response to infection by *Mycobacterium tuberculosis*. The adaptive immune system of the adult stage zebrafish offers an advance on the commonly used embryo infection model as adult zebrafish form granulomas with striking similarities to human-*M*. *tuberculosis* granulomas. Here, we present workflows to perform high content analyses of granulomas in adult zebrafish infected with *M. marinum* by cryosectioning to take advantage of strong endogenous transgenic fluorescence adapted from common zebrafish embryo infection tools. Specific guides to classifying granuloma necrosis and organisation, quantifying bacterial burden and leukocyte infiltration of granulomas, and visualizing extracellular matrix remodelling and foam cell formation are also provided. We use these methods to characterize neutrophil recruitment to *M. marinum* granulomas across time and find an inverse relation to granuloma necrosis suggesting granuloma necrosis is not a marker of immunopathology in the natural infection system of the adult zebrafish-*M*. *marinum* pairing. The methods can be easily translated to studying the zebrafish adaptive immune response to other chronic and granuloma-forming pathogens.

## 1. Introduction

Granulomas are the structural hallmark of human tuberculosis caused by infection with *Mycobacterium tuberculosis*. Tuberculous granulomas are comprised of host immune cells from the innate and adaptive lineages that act in concert to physically contain mycobacteria. Recent research has provided evidence that pathogenic mycobacteria actively drive granuloma formation, and associated inflammation, in a strategy to evade immune control (Ramakrishnan, 2012).

The *zebrafish-Mycobacterium marinum* infection system is an important model of human tuberculosis. Transparent zebrafish embryos have been extensively utilized to understand the early pathogenesis of mycobacterial infection and specifically the host-microbe interactions mediated by innate immune cells. The innate immune cell granulomas in zebrafish embryos recapitulate several important aspects of human-*M*. *tuberculosis* granulomas including macrophage epithelioid differentiation, necrosis, and the dynamic recruitment of susceptible naïve macrophages and egress of infected macrophages (Davis et al., 2002; Davis and Ramakrishnan, 2009). However, zebrafish embryos lack an adaptive immune system and generally lose control of infection within 5-7 days precluding any investigation of sterilizing or latent granulomas.

Infection of adult zebrafish with *M. marinum* provides the addition of the adaptive immune system to the zebrafish-*M*. *marinum* platform. Adult zebrafish form granulomas that are structurally closer to the human*-Mycobacterium tuberculosis* granuloma than that seen in inbred mouse-*M*. *tuberculosis* granulomas, and recapitulate important aspects of human granulomas such as hypoxia, sterilizing immunity and latency (Oehlers et al., 2015; Parikka et al., 2012). Histological examination of the adult zebrafish-*M*. *marinum* infection system has been instrumental in advancing our understanding of mycobacterial virulence, granuloma macrophage epithelioid differentiation, vascularization, and the adaptive immune response to mycobacterial infection (Cronan et al., 2016; Oehlers et al., 2015; Parikka et al., 2012; Prouty et al., 2003; Risalde et al., 2018; Swaim et al., 2006; van der Sar et al., 2004; Volkman et al., 2004; Weerdenburg et al., 2012).

Here, we present our simple methodology for generating specimens and analysis of granulomas from cryosection-generated slides. We almost exclusively utilize frozen rather than paraffin sectioning to take advantage of the strong native fluorescence afforded by *msp12* promoter-driven fluorescence in *M*. *marinum* (pTEC plasmids available https://www.addgene.org/Lalita_Ramakrishnan/) and the wide range of immune cell lineage transgenic markers available in zebrafish (see Table 1) (Takaki et al., 2012; Takaki et al., 2013).

**Table 1:** Zebrafish reporter lines and visualisation strategies

| Line | Cell type(s) marked | Visualisation strategy | Reference |
| --- | --- | --- | --- |
| Tg(cd41:GFP<sup>la2</sup>) | Thrombocytes | Native fluorescence or anti-GFP boost | (Lin et al., 2005) |
| Tg(kdrl:GFP<sup>s843</sup>) | Endothelial cells | Native fluorescence | (Jin et al., 2005) |
| Tg(lyzC:dsRed<sup>nz50</sup> or GFP<sup>nz117</sup>) | Neutrophils | Native fluorescence | (Hall et al., 2007) |
| Tg(mfap4:iCre-2A-Tomato<sup>xt8</sup>, ubb:LOXP-TagBFP-LOXP-Tomato<sup>xt7</sup>)) | Macrophages | Native fluorescence | (Cronan et al., 2016) |
| TgBAC(foxp3a:TagRFP<sup>vcc6</sup>) | T regulatory cells | Native fluorescence | (Hui et al., 2017) |
| TgBAC(lck:EGFP<sup>vcc4</sup>) | T cells | Anti-GFP boost | (Sugimoto et al., 2017) |
| TgBAC(pdgfrb:gfp<sup>ncv22</sup>) | Fibroblasts and perivascular cells | Native fluorescence or anti-GFP boost | (Ando et al., 2016) |
| TgBAC(tnfa:GFP<sup>pd1028</sup>) | Tumor necrosis factor expressing cells | Native fluorescence | (Marjoram et al., 2015) |

## 2. Critical Experimental Materials

Water for fish: 1 g/L sea salt water or aquarium system water.

Dry fish food: we use O.range GROW (INVE Aquaculture). Similar results are expected with any solid pellet or flake shaped food where it is simple to remove uneaten debris. We have avoided the use of live feeds due to difficulty in cleaning uneaten feed from beakers.

Fluorescent *M*. *marinum*: We utilize an abbreviated version of the method described by Takaki *et al*., bacteria are grown in 7H9 liquid media supplemented with OADC (Sigma-Aldrich M0678), Tween-80 (Sigma-Aldrich P6474, final concentration 0.0045%) and 50 g/l hygromycin to a mid log phase (OD600 ~0.6-0.8). Bacteria are harvested by repeated centrifugation, passage through 29 G needles, and resuspension in 7H9 supplemented with OADC only to prepare a single cell suspension. This single cell suspension is then frozen at −80°C and diluted as necessary for infection experiments (Takaki et al., 2013).

PBS: Phosphate buffered saline.

Phenol red dye: 0.5% in Dulbecco’s Phosphate Buffered Saline, sterile-filtered (Sigma-Aldrich P0290).

Parafilm.

Tricaine: 15x tricaine stock is 4 g/l MS-222 (Ethyl 3-aminobenzoate methanesulfonate, Sigma-Aldrich E10521) dissolved in deionized water and pH adjusted to 7 with sodium hydroxide.

Injection needles: 31 G BD Ultra-Fine II Short Needle 0.3 ml syringe. Provides similar results to Hamilton Syringes with the ease of disposable price.

Bleach: final concentration of decontaminating solution should be 1% available bleach. Generally this is achieved with a 10% volume of commercially available 10% bleach poured into a carboy prior to addition of contaminated liquids.

Fixative: 10% neutral buffered formalin or 16% paraformaldehyde for dilution into media at a 1:3 ratio for a final concentration of 4% paraformaldehyde.

30% (w/v) sucrose: 30 g sucrose / 100 ml deionized water, filter sterilized.

OCT: Optimal Cutting Temperature (OCT) compound. We currently use Sakura #4583 but have used a variety of commercial suppliers and have not observed appreciable differences.

High quality adhesion microscope slides: We currently use SuperFrost Ultra Plus© Thermo Scientific. Lower grades of slides have been more prone to tissue loss.

Blocking solution: The majority of our secondary antibodies were raised in goat so we routinely use normal goat serum diluted to 5% in PBS.

Primary antibody to boost GFP signal: Chicken Anti-GFP (Abcam: ab13970)

Secondary antibody to boost GFP signal: Goat anti-Chicken IgY (H+L), Alexa Fluor^®^ 488 (Abcam: ab150169)

Fluoromount with DAPI: We currently use DAPI Fluoromount (Proscitech IM035) but have achieved similar results with anti-fade mounting media with DAPI from a range of suppliers. FIJI-modified ImageJ: Free download at https://fiji.sc/ allows simple opening of proprietary microscope image formats and enhanced functionality on top of ImageJ (Schindelin et al., 2012; Schneider et al., 2012).

## 3. Infection procedure

1. Collect zebrafish from your aquarium system into clean water for fish within 1000 ml beakers covered by tin foil. We typically house 5-6 zebrafish in 500 ml liquid volume within a 1000 ml beaker. Change ~25% water daily to remove fecal matter and feed with dry fish food.
2. Acclimatize zebrafish to new housing for period determined by your animal welfare code. Beakers should be kept in 28°C environment with light cycle matching your aquarium.
3. Set up injection station with equipment illustrated in Figure 1.
4. Thaw aliquots of *M. marinum* and dilute in sterile PBS and phenol red dye to produce a ~200CFU concentration of bacteria per 10 or 15 μl. Typically we perform the final dilution into 450 μl PBS and 50 μl phenol red dye.
5. Spot 10-15 μl of inoculum onto parafilm so that there is one spot per zebrafish plus a “spare” spot to be ready to compensate for mistakes. We generally inject 15 μl into cohorts of animals larger than 30 mm Standard Length.
6. Anesthetize zebrafish in 0.6-0.75x tricaine in groups of up to 5-6 depending on user speed. We typically start training users by anesthetizing 2 zebrafish at a time.
7. Draw up bacterial inoculum into injection needle with bevel down while zebrafish lose consciousness.
8. Transfer a single zebrafish onto wetted sponge and position with ventral side towards your dominant hand.
9. Use a finger on your non-dominant hand to secure the zebrafish against the wetted sponge and inject into the anaesthetized adult fish under the “armpit” of the pelvic fin into the intraperitoneal space holding the injection needle and syringe in your dominant hand. Correctly injected fish will have red dye visible within the peritoneal cavity. Common incorrect injections can result in significant dye leakage out (incorrect angle of injection) or subdermal spread of dye (insufficient penetration through the skin).
10. Recover injected fish back into clean water for fish at 28°C and monitor for recovery from anesthetic. Animals typically recover within 2-5 minutes from the low dose of anesthetic used in this procedure.
11. House infected zebrafish in a dedicated 28°C incubator fitted with a light cycle matching your aquarium. A simple household power outlet timer and an LED lamp fixed to a standard air jacket incubator is a cost-effective solution to maintain containment of BSL2 *M. marinum*-infected zebrafish.
12. Monitor infected fish daily. We typically house 5-6 zebrafish in 500 ml liquid volume within a 1000 ml beaker. Fecal matter and 200-250 ml water are removed by pipetting with a 50 ml pipette and decontaminated in bleach. Water is replaced and animals are fed standard dry fish food. Take care to only feed as much food as is rapidly consumed, the small water volumes are highly susceptible to spoiling by rotting food. Zebrafish will typically lose their appetite in the first 2-3 days post infection.
13. Euthanize zebrafish from 2 weeks post infection to observe stereotypical granulomas. We utilize tricaine overdose although comparable results are expected for other methods of euthanasia such as ice water bath.

**Figure 1:**
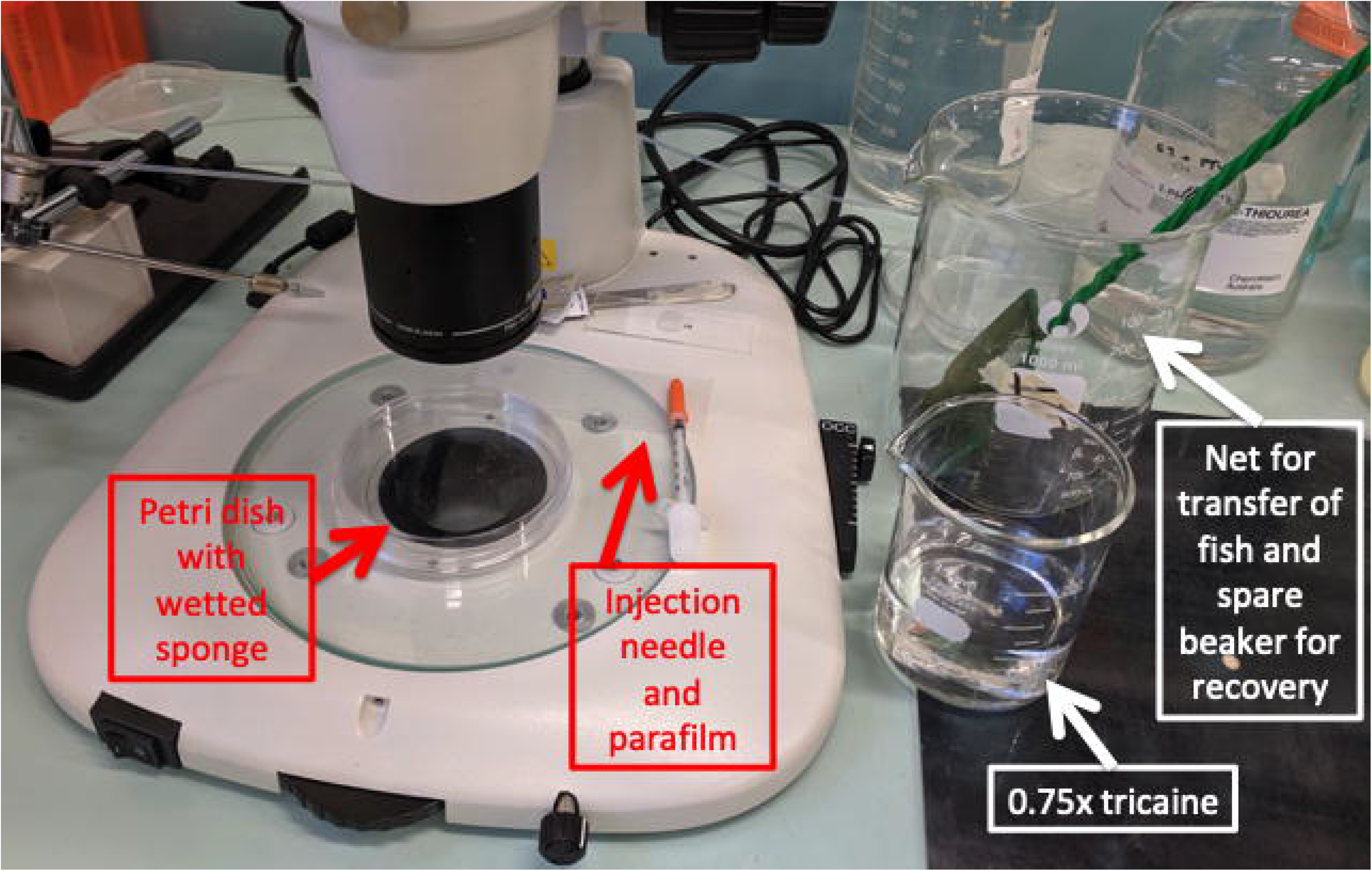
Microscope station set up for intraperitoneal injection of *M. marinum* into adult zebrafish. Injection station set up for a right-handed operator.

## 4. Preparation for cryosectioning

1. Transfer euthanized zebrafish to Petri dish.
2. Use two tweezers to guillotine the tail off at a point caudal to the cloaca. This will aid fitting the zebrafish into a cryomold.
3. Use tweezers perform an incision into the side of the peritoneal cavity skin taking care not to disrupt the internal organs.
4. Optional: decapitate zebrafish using two tweezers to guillotine the head off at the gills. Although observed by other groups using the *M. marinum* E11 strain, we have never observed *M. marinum* M strain dissemination to the head following intraperitoneal injection (van Leeuwen et al., 2014). Decapitation may also aid fitting particularly large zebrafish into a cryomold.
5. Fix in fixative for 1-3 days at 4 °C. Longer periods may diminish endogenous fluorescence.
6. Wash fixed specimens in PBS for at least 1 hour at room temperature.
7. Optional: decalcify in 0.5M EDTA. This step is not necessary in our hands as microtome blades easily cut through bone and scales.
8. Replace PBS with 30% (w/v) sucrose solution overnight at room temperature.
9. Remove 50% volume of sucrose and replace with OCT compound for a 50:50 final ratio and incubate overnight at room temperature.
10. Replace mixture with OCT compound and incubate overnight at room temperature.
11. Transfer specimens to cyromolds and cover with OCT compound.
12. Freeze embedded specimens in −80 °C freezer for at least 1 hour.

## 5. Cryosectioning, fluorescence staining and imaging

1. Prewarm specimens and cool microtome blade to −20 °C in cryostat.
2. Mount specimen, trim as necessary, and align parallel to microtome blade.
3. Section at 20 μm and mount onto high quality adhesion microscope slides. We only collect sections once the peritoneal cavity is visible as we have not observed mycobacterial dissemination to the skin or muscle.
4. Optional: to save time and reduce redundancy between adjacent sections, we typically collect 3 sections onto an “A” slide, the next 3 sections onto a “B” slide and then discard the next 4 sections resulting in slides containing sections spaced 200 μm apart. We routinely collect 6-8 sections per standard microscope slide yielding approximately 10-20 slides per infected zebrafish.
5. Label slides and store in at −20°C in cryostat until sectioning is complete.
6. Optional: use a fluorescent dissecting microscope to check slides for fluorescent bacteria. Discard early and late slides that do not contain fluorescent bacteria. Do not overexpose slides at this point, fluorescence bleaches rapidly in OCT.
7. Store slides at −80°C until imaging.
8. Thaw slides for 2-5 minutes at room temperature.
9. Wash slides 1 or 2 times in PBS for 5 minutes to remove OCT. Rinsing is critical as residual OCT interferes with downstream fluorescence microscopy.
10. Optional antibody staining: determine if your fluorophores of interest require signal boosting with fluorescently labeled antibodies (Table: Zebrafish reporter lines and visualization strategies) or if you are combining native transgenic fluorescence with additional antibody detection targets (such as hypoxyprobe, E-cadherin, L-plastin).

a. Postfix slides in fixative for 1-2 minutes at room temperature. Although longer incubations will preserve sections during subsequent wash steps, refixing diminishes native fluorescence.
b. Rinse slides twice in PBS for 5 minutes to remove fixative.
c. Block slides for 1 hours at RT with blocking solution of choice by gentle pipetting onto the top of the slide. Cover with parafilm.
d. Flick off blocking solution and gently pipette 100-150 μl diluted primary antibody onto the slide. Cover with parafilm, add water to slide box to humidify, and incubate at 4°C overnight.
e. Next day, rinse slides with 3-4 changes of PBS over 30 minutes to remove unbound primary antibody.
f. Gently pipette 100-150 μl diluted secondary fluorescent antibody onto the slide. Cover with parafilm, add water to slide box to humidify, and incubate at 4°C overnight or 3-4 hours at room temperature.
g. Rinse slides with 3-4 changes of PBS over 30 minutes to remove unbound secondary antibody. Notes: sections can be highly susceptible to damage during washing and antibody addition steps. Take care to reduce sheer forces when moving slides and pipetting.
11. Apply 1-3 drops of fluoromount with DAPI and cover slide with coverslip.
12. Image on microscope of choice collecting each channel as a separate file or in a format that allows splitting of channels in ImageJ. We utilize microscopes with a “stitch” or “mosaic” feature which allows the assembly of a whole section frame of view from several individual fields of view.

## 6. Fluorescence image analysis

1. Open file(s) in FIJI-modified ImageJ. If necessary, split image into individual channels using the menu item Image>Color>Split Channels.
2. Adjust the brightness parameters of each channel to minimize background fluorescence and maximize positive signal using the menu item Image>Adjust>Brightness/Contrast. Optimizing the DAPI and bacterial fluorescence channels will assist in accurately determining the edges of granulomas in subsequent steps.
3. Merge at least your DAPI and bacterial fluorescence channels into a recolorized image using the menu item Image>Color>Merge Channels. Activate “Create composite” and “Keep source images”, ensure “Ignore source LUTs” is deactivated.
4. Use the “Freehand selections” tool to trace the edges of a granuloma (Figure 2A). Additional channels of immune cells may aid identification of granuloma edges, or potentially bias analysis.
5. Press the “t” button on your keyboard to add the selection to your region of interest (ROI) manager list.
6. Repeat steps 4 and 5 until all granulomas are assigned a ROI.
7. Optional: save the ROI list using the ROI Manager menu command More>Save. This is useful if you are going to perform image analyses across different sessions.
8. Optional: save the overlay image for reference.
9. Quantify bacterial burden.

a. Reopen the bacterial channel image or revert from the brightness-adjusted image using the menu item File>Revert.
b. Record a macro with the following instructions, macro code lines are *italicized* and available in the Supplementary data:

i. Convert file to 8-bit: Image>Type>8-bit= *run(“8-bit”J;*
ii. Remove scale to produce results in pixels: Analyze>Set Scale>“Click to Remove Scale”, “OK” = *run(“Set Scale…”, “distance=0”);*
iii. Set thresholding to highlight pixels above threshold = *setAutoThreshold(“Default dark”);*
iv. Set thresholding limits to lowest specific bacterial fluorescence intensity as “XX” and 255 as maximal signal: Image>Adjust>Threshold>“Set” = *setThreshold(XX, 255);*
v. Either manual mode Count pixels above threshold: Analyze>Analyze Particles>Check “Summarize” = *run(“Analyze Particles…”, “summarize”);*
vi. Or automatic mode Repeat operation through ROI list=

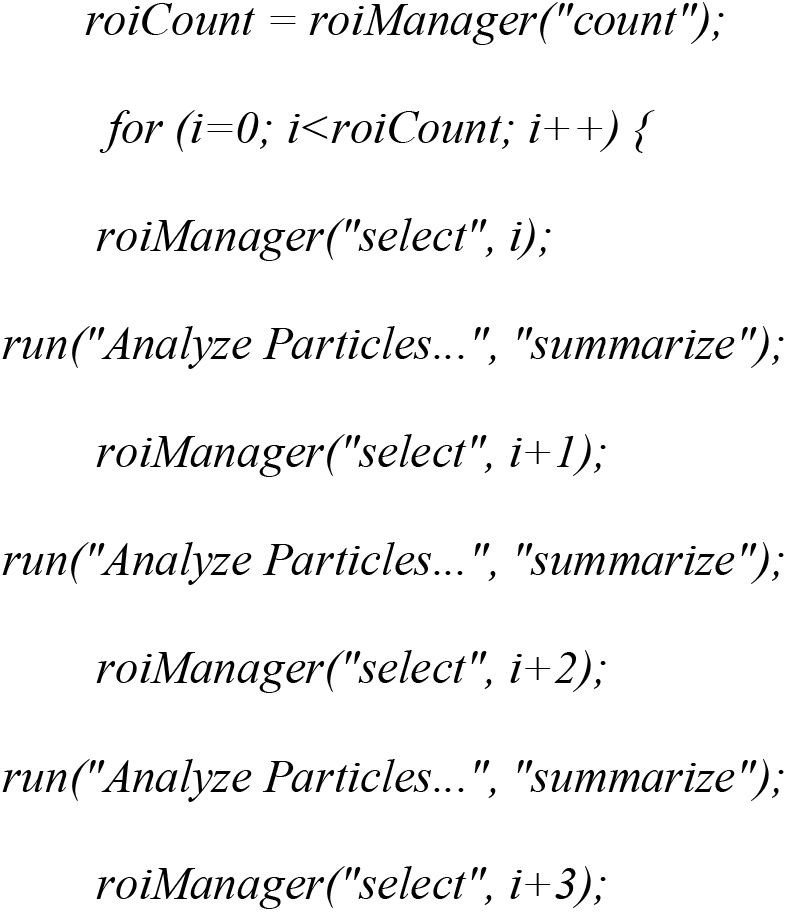 Command can be repeated by duplicating the repeating two lines and continuing the sequence i+1, i+2, i+3, i+4 to accommodate the size of your ROI list. The Supplementary data for this paper contains a macro to count up to 70 regions of interest.
vii. Run the macro.
viii. Copy the Summary window data to your spreadsheet software of choice. The “Total Area” column provides bacterial burden per user-defined granuloma.
10. Classify granuloma morphology from the DAPI channel (Cronan et al., 2016).

a. Cellular vs necrotic

i. Switch to the brightness-adjusted DAPI channel image.
ii. Check the “Show All” and “Labels” boxes in the ROI Manager window to highlight all user-defined granulomas.
iii. Score each granuloma in sequence for the presence of central necrosis (Figure 2B).
b. Epithelialized vs disorganized

i. Switch to the brightness-adjusted DAPI channel image.
ii. Check the “Show All” and “Labels” boxes in the ROI Manager window to highlight all user-defined granulomas.
iii. Score each granuloma for directional organization of host nuclei at the rim of the granuloma (Figure 2B).
11. Quantify interaction of reporter marked cells with granulomas.

a. Leukocyte infiltration as an example of discrete data.

i. Reopen the leukocyte channel image or revert from the brightness-adjusted image using the menu item File>Revert. If necessary, reload the saved ROI list.
ii. Open the macro created in Step 9b to quantify bacterial burden. Adjust the lower limit of the threshold “XX” in Step 9b-iv to lowest specific leukocyte fluorescence intensity that removes background signal.
iii. Run the macro.
iv. Copy the Summary window data to your spreadsheet software of choice. The “Total Area” column provides leukocyte pixel units per user-defined granuloma. Leukocyte pixel units can be used as an arbitrary measurement of leukocyte number or converted to Leukocyte units by calibrating to the pixel area of single discrete leukocytes (Ellett and Lieschke, 2012). We do not believe the “Count” column in the Summary window provides an accurate estimation of leukocyte numbers as leukocytes are relatively amorphous compared to round colonies.
b. Blood vessel proximity as an example of continuous data (Oehlers et al., 2015).

i. Create a new two channel overlay of the brightness-adjusted bacterial and blood vessel channels using the menu item Image>Color>Merge Channels. Activate “Create composite” and “Keep source images”, ensure “Ignore source LUTs” is deactivated.
ii. If necessary, reload the saved ROI list. Check the “Show All” and “Labels” boxes in the ROI Manager window to highlight all user-defined granulomas.
iii. Remove scale to produce results in pixels: Analyze>Set Scale>“Click to Remove Scale”, “OK”
iv. Use the “Straight line tool” to draw the shortest distance between the edge of bacterial fluorescence and the nearest blood vessel.
v. Use the menu command Analyze>Measure to calculate the length of the user-drawn straight line (the “Length” column of the Results window).
vi. Repeat Steps iv. to v. in ROI list sequence until all granulomas are assigned a minimum distance from vasculature.
vii. Export data to spreadsheet of choice to correlate with other granuloma parameters. Convert pixel units to μm using appropriate conversion factor.

**Figure 2:**
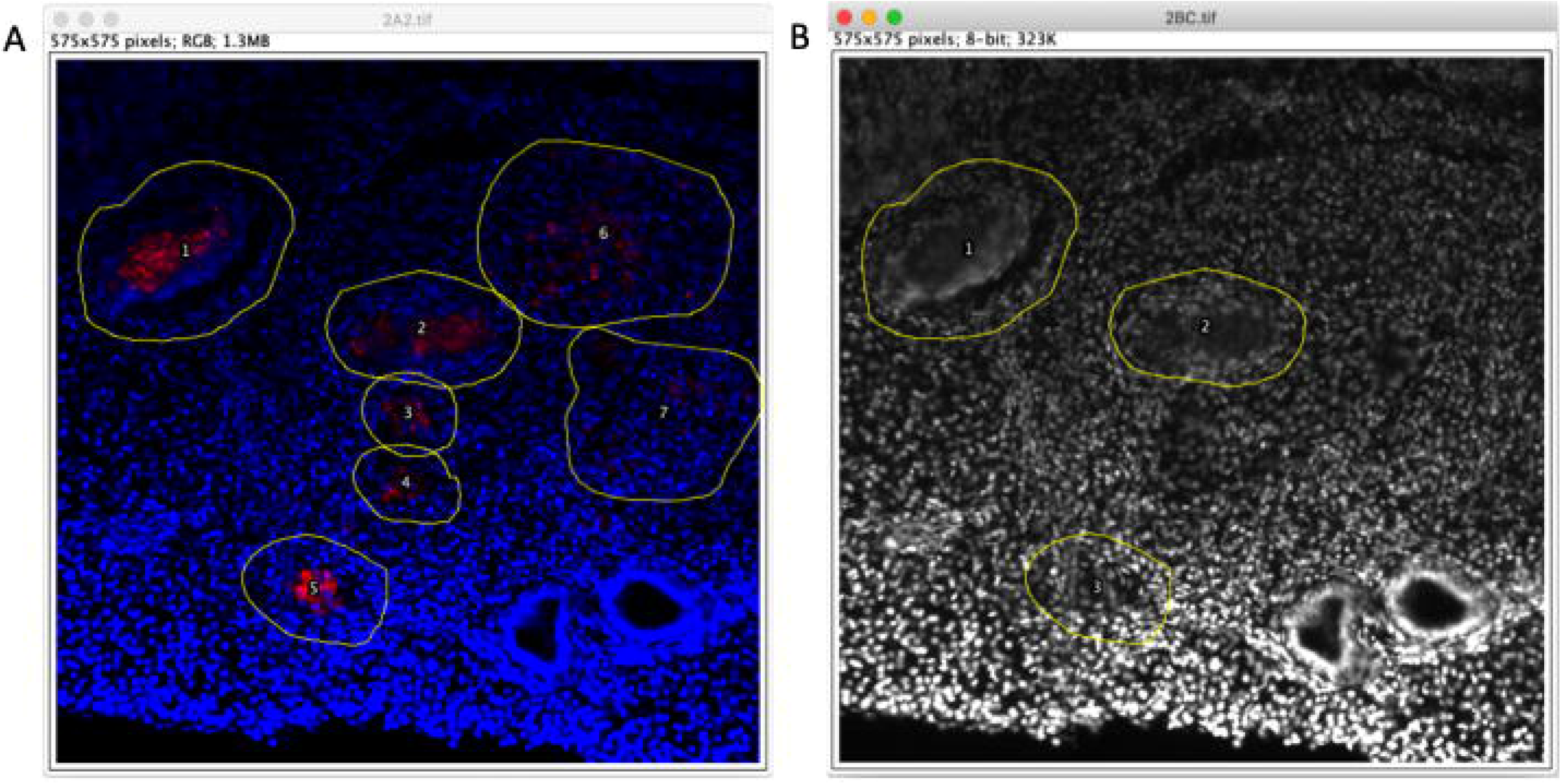
Defining and categorizing granuloma morphology. (A) Use of two color overlay and the “Freehand selections” tool to trace the edges of a granuloma and annotate as number regions of interest (ROIs) (B) Use of DAPI channel to identify the presence of central necrosis highlighted in ROIs 1 and 2, and directional organization of host nuclei at the rim of the granuloma highlighted in ROIs 1, 2, and 3 (previously annotated as #5 in (A)).

## 7. Non-fluorescence correlative staining and microscopy

This section takes advantage of the library of adjacent slides created by the A/B section spacing described in step 5.4 to correlate bacterial distribution characterized in step 6 with histological stains that destroy native fluorescence or are not compatible with fluorescence microscopy. This section continues from step 5.9.

1. Oil Red O staining for foam cells (protocol for isopropanol solvent is similar just substitute isopropanol of propylene glycol in steps c, d, e) (Johansen et al., 2018).

a. Postfix slides in fixative for 5-10 minutes at room temperature.
b. Filter 0.5% (w/v) Oil Red O (Sigma-Aldrich O0625) dissolved in propylene glycol to remove precipitate.
c. Rinse slides twice in PBS for 5 minutes to remove fixative.
d. Rinse slides twice in propylene glycol for 5 minutes.
e. Stain slides in 0.5% (w/v) Oil Red O propylene glycol solution for 15 minutes.
f. Rinse slides twice in propylene glycol for 5 minutes to remove background staining.
g. Rinse slides briefly in PBS.
h. Counterstain slides with a 1% (w/v) solution of methylene blue (Sigma-Aldrich M9140) dissolved in water or hematoxylin for 1 minute.
i. Rinse slides briefly in tap water.
j. Add 2-3 drops of with aqueous mounting media (Clear-Mount, Proscitech IM032) and mount coverslip.
k. Image by light microscopy.
2. Picrosirius red staining for extracellular matrix remodeling.

a. Postfix slides in fixative for 5-10 minutes at room temperature.
b. Rinse slides briefly twice in tap water to remove fixative.
c. Cover slides in picrosirius red (Polysciences #24901 or 0.1% (w/v) Sirius red F3B dissolved in saturated picric acid) and incubate for 1 hour at room temperature.
d. Rinse slides briefly twice in 0.1 N hydrochloric acid.
e. Rinse slides in tap water.
f. Dehydrate slides in 70% ethanol.
g. Add 2-3 drops of ethanol-compatible mounting media (Entellan, Sigma-Aldrich 107960) and mount coverslip.
h. Image by light or polarized light microscopy (Figure 3).

**Figure 3:**
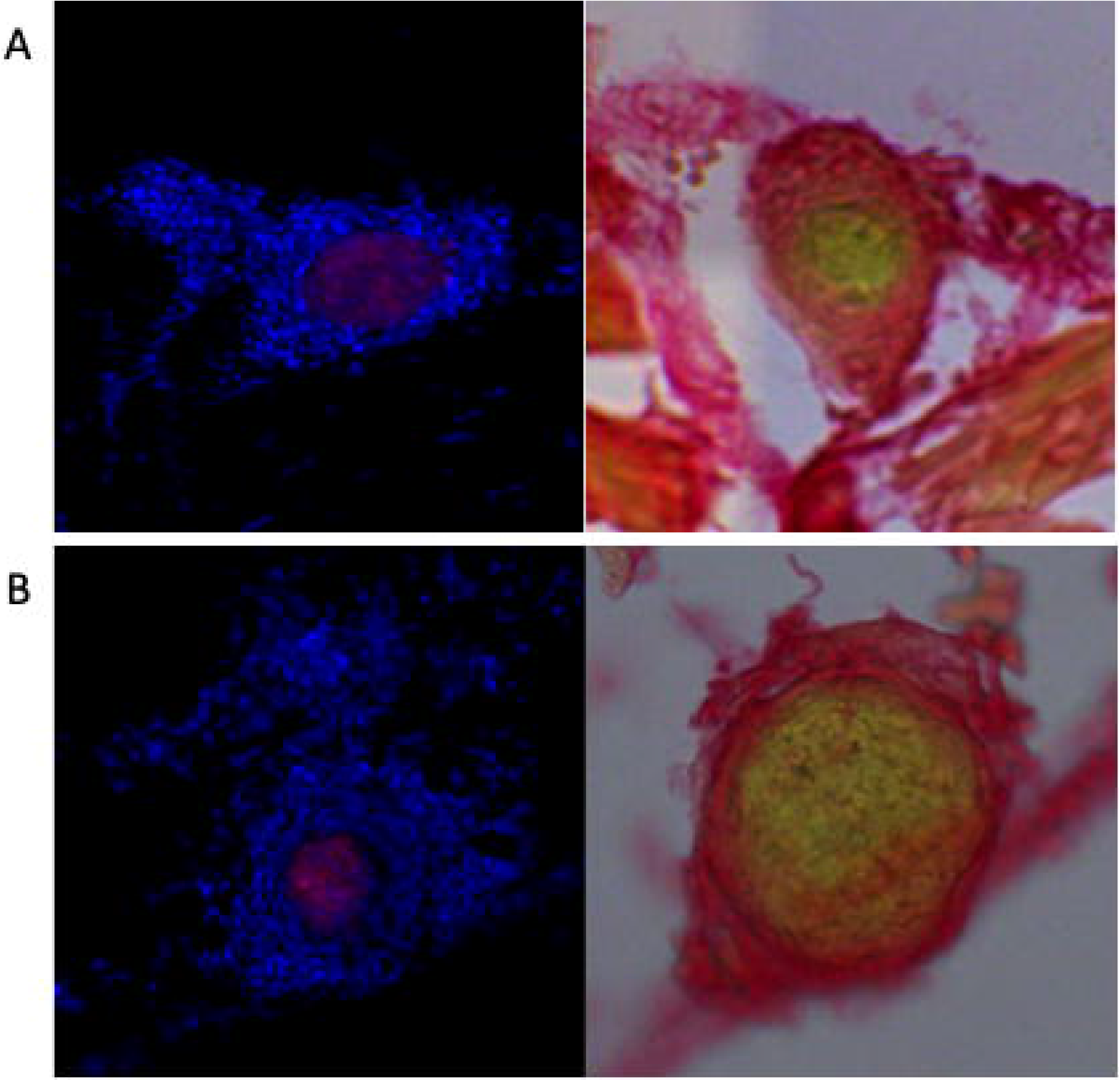
Extracellular matrix reorganization around zebrafish-*M*. *marinum* granulomas visualized by picrosirius red staining. (A) Matched DAPI and *M*. *marinum*-tdTomato fluorescence image and picrosirius red-stained bright field image of a granuloma in the head kidney demonstrating collagen (red) deposition through the cellular body of the necrotic granuloma. (B) Matched DAPI and *M*. *marinum*-tdTomato fluorescence image and picrosirius red-stained bright field image of a granuloma in the liver demonstrating highly organized collagen (red) containment of a necrotic granuloma.

## 8. Characterization of neutrophil recruitment to granulomas

We applied the methods described in Sections 3-6 on *Tg(lyzC:DsRed^nz50^)* zebrafish infected with *M. marinum*-wasabi to characterize neutrophil infiltration of granulomas across time, and as a function of bacterial burden and granuloma morphology (Figure 4A). Two to three animals were harvested at each of 1, 2, 4, and 6 weeks post infection and analyzed by our census technique generating 53, 81, 24, and 125 granulomas at each timepoint respectively.

**Figure 4:**
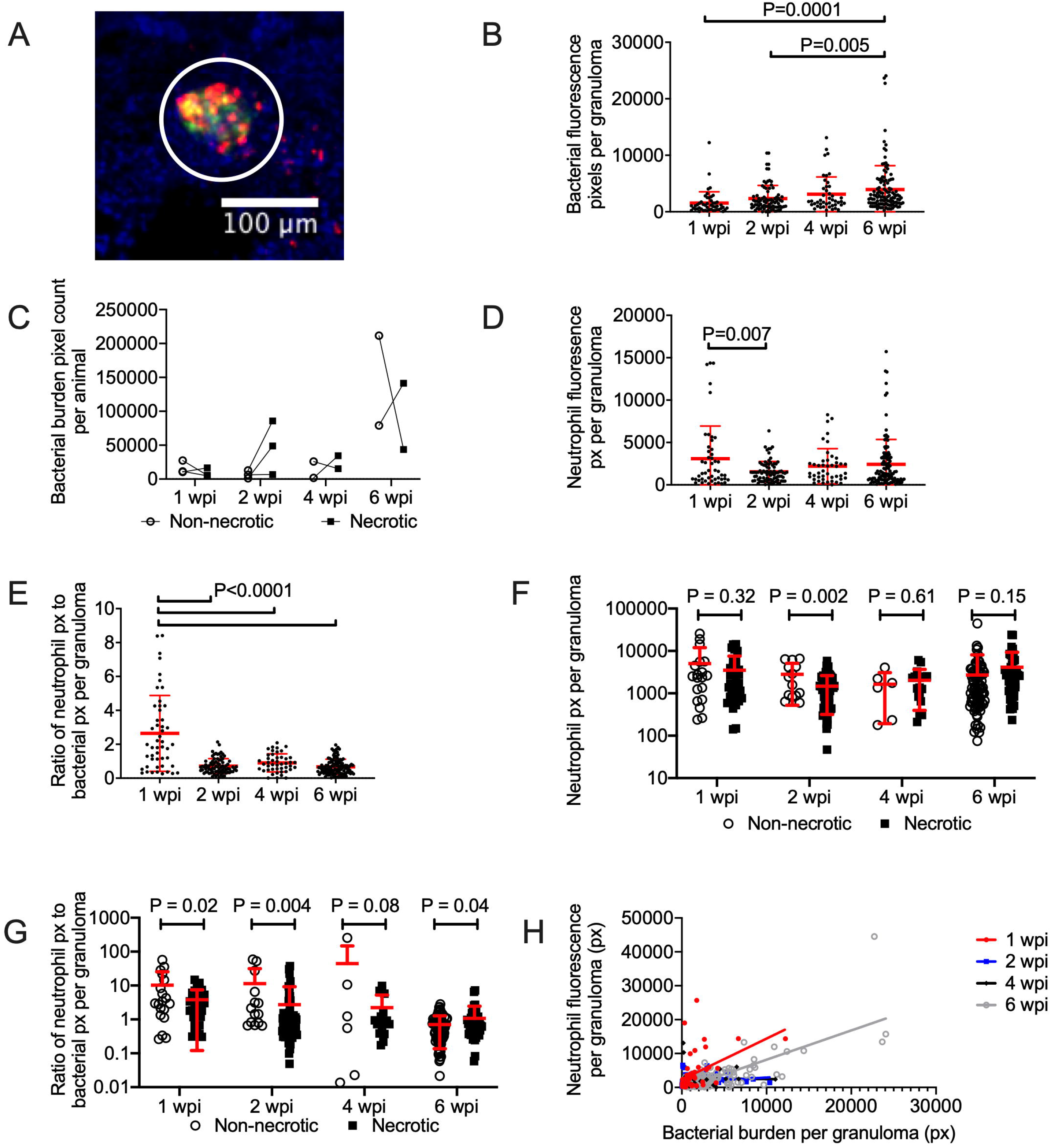
Analysis of neutrophil recruitment to granulomas in the adult zebrafish-*M*. *marinum* infection model. (A) Representative image of granuloma from a DAPI-stained section from a *Tg*(*lyzC:dsred*)^*nz50*^ adult zebrafish infected with *M*. *marium*-wasabi. White circle indicates edge of granuloma as defined by inspection of DAPI channel covered in Section 6.4. (B) Quantification of granuloma bacterial content by fluorescent pixel count in individual granulomas. (C) Quantification of total bacterial content per animal stratified by the absence or presence of necrosis in each lesion. P values from matched T-tests performed for each timepoint. (D) Quantification of neutrophil recruitment to granulomas by fluorescent pixel count in individual granulomas. (E) Ratio of neutrophil fluorescent pixels divided by bacterial fluorescent pixels in individual granuloma. (F) Quantification of neutrophil recruitment to granulomas by fluorescent pixel count in individual granulomas stratified by stratified by the absence or presence of necrosis in each lesion. (G) Ratio of neutrophil to bacterial fluorescent pixels in individual granulomas stratified by stratified by the absence or presence of necrosis in each lesion. (H) Linear regression analysis of neutrophil and bacterial fluorescent pixels in individual granulomas. P values from one way ANOVA with Tukey multiple comparison unless otherwise indicated.

As expected from previous reports of zebrafish-*M*. *marinum* granuloma coalescence (Cronan et al., 2016; Cronan et al., 2018; Oehlers et al., 2015), bacterial burden per granuloma increased over time (Figure 4B). Our high content analysis further allowed us to stratify granulomas into non-necrotic (cellular) or necrotic categories. Total bacterial burden was evenly distributed across necrotic compared to non-necrotic granulomas at all timepoints with only a trend to a higher total of burden within necrotic granulomas seen at 2 wpi (P=0.16) illustrating the heterogeneity of granuloma classes in the zebrafish model (Figure 4C).

Analysis of neutrophil infiltration across time revealed fairly stable neutrophil recruitment per granuloma with a slight reduction at 2 wpi correlating with appearance of visible granuloma organization, neutrophil recruitment then trended toward gradually increasing later in infection at 4 and 6 wpi (Figure 4D). The ratio of neutrophils to bacteria per granuloma peaked at 1 wpi followed by a significantly lower ratio across 2, 4 and 6 wpi, suggesting a more controlled inflammatory response following the initial spike at 1 wpi (Figure 4E). Stratification of granulomas by necrotic status revealed reduced neutrophil recruitment to necrotic granulomas at 2 wpi, but not the other timepoints (Figure 4F). This observation was accentuated by normalizing for bacterial burden in each granuloma with significantly fewer neutrophils around necrotic granulomas for the first two weeks of infection (Figure 4G).

Interestingly, linear regression analysis found only a weak relationship between neutrophil infiltration and bacterial burden per granuloma at 1 wpi (R square 0.20, non-zero slope P=0.0007), no relationship at 2 and 4 wpi, and a strong relationship at 6 wpi (R square 0.57, non-zero slope P<0.0001) suggesting an early burden-dependent influx of neutrophils that lessens during granuloma maturation and organization until later in infection when granulomas break down restarting the cycle (Figure 4H).

## 9. Conclusions

The methodologies presented here provide a blueprint to perform high content analyses of adult zebrafish-*M*. *marinum* granulomas. Our suggested census approach using correlative acquisition of quantitative data seeks to reproduce two important aspects of the zebrafish embryo-*M*. *marinum* infection model: *in toto* analysis of infection and high specimen numbers providing statistical power.

Advances in optical clearing of whole zebrafish for 3-dimensional *in situ* analysis of granulomas is a powerful tool for relatively artifact-free microscopy but requires access to expensive multiphoton microscopes to access deep tissues (Cronan et al., 2015). Our method attempts to census the granuloma load of individual zebrafish by creating a catalog of thin section snapshots at 200 μm intervals providing confidence that granulomas are not double counted. The use of explanted adult zebrafish-*M*. *marinum* granulomas is expected to provide insight into the 4-dimensional behavior of granulomas by serial live imaging and could be used to functionally investigate the correlative datasets produced by our methodology (Cronan et al., 2018).

Our analysis of bacterial burden in granuloma classes and neutrophil recruitment to granulomas over the course of intraperitoneal infection revealed two important findings. Firstly, the 2 wpi timepoint is distinct from earlier and later timepoints. Complementing our previous work demonstrating increased organization of granulomas at the 2 wpi timepoint (Cronan et al., 2016; Oehlers et al., 2015), we have documented both a trend to most bacteria being located within organized necrotic granulomas and a reduction in neutrophil infiltration to these necrotic granulomas at this timepoint. The initial peak in neutrophil recruitment to sites of *M*. *marinum* at 1 wpi is a clear correlate with the initial inflammatory response seen in mammalian models of *M*. *tuberculosis* infection.

Secondly, our data suggest granuloma necrosis may be a natural step in the effective control of mycobacterial infection in the zebrafish model. Neutrophilic inflammation is a marker of granuloma-associated immunopathology in mammals and is usually associated with necrotic breakdown of granulomas (Barnes et al., 1988; Eruslanov et al., 2005; Eum et al., 2010). Conversely, our data shows reduced neutrophilic recruitment to necrotic granulomas compared to cellular lesions during early, presumably mostly primary, infection suggesting the formation of organized necrotic granulomas are the outcome of balancing immune control of infection without significant immunopathology in the adult zebrafish-*M*. *marinum* infection model. Previous work has shown granuloma macrophage epithelization excludes neutrophils from accessing granulomas in the zebrafish providing a mechanism for our observation (Cronan et al., 2016).

Zebrafish are susceptible to a range of chronic granuloma-forming infections by important human pathogens such as *Cryptococcus neoformans* and *Mycobacterium abscessus* (Bernut et al., 2014; Tenor et al., 2015). Our methodology can be easily translated to studying the zebrafish immune response to these pathogens with only minor changes to infectious dose in order to provide additional comparative models of host-pathogen interactions.

## Supporting information

Macro template

## Acknowledgements

We thank Drs Mark Cronan and David Tobin at Duke University School of Medicine, and members of the Oehlers lab at the Centenary Institute for helpful discussion of techniques and troubleshooting.

This work was supported by an Australian National Health and Medical Research Council CJ Martin Early Career Fellowship APP1053407 and Project Grant APP1099912; The University of Sydney Fellowship G197581; NSW Ministry of Health under the NSW Health Early-Mid Career Fellowships Scheme H18/31086; and the Kenyon Family Foundation Inflammation Award (S.H.O.).

## References

Ando, K., Fukuhara, S., Izumi, N., Nakajima, H., Fukui, H., Kelsh, R.N., Mochizuki, N., 2016. Clarification of mural cell coverage of vascular endothelial cells by live imaging of zebrafish. Development 143, 1328–1339.

Barnes, P.F., Leedom, J.M., Chan, L.S., Wong, S.F., Shah, J., Vachon, L.A., Overturf, G.D., Modlin, R.L., 1988. Predictors of short-term prognosis in patients with pulmonary tuberculosis. J Infect Dis 158, 366–371.

Bernut, A., Herrmann, J.L., Kissa, K., Dubremetz, J.F., Gaillard, J.L., Lutfalla, G., Kremer, L., 2014. Mycobacterium abscessus cording prevents phagocytosis and promotes abscess formation. Proc Natl Acad Sci U S A 111, E943–952.

Cronan, M.R., Beerman, R.W., Rosenberg, A.F., Saelens, J.W., Johnson, M.G., Oehlers, S.H., Sisk, D.M., Jurcic Smith, K.L., Medvitz, N.A., Miller, S.E., Trinh, L.A., Fraser, S.E., Madden, J.F., Turner, J., Stout, J.E., Lee, S., Tobin, D.M., 2016. Macrophage Epithelial Reprogramming Underlies Mycobacterial Granuloma Formation and Promotes Infection. Immunity 45, 861–876.

Cronan, M.R., Matty, M.A., Rosenberg, A.F., Blanc, L., Pyle, C.J., Espenschied, S.T., Rawls, J.F., Dartois, V., Tobin, D.M., 2018. An explant technique for high-resolution imaging and manipulation of mycobacterial granulomas. Nat Methods 15, 1098–1107.

Cronan, M.R., Rosenberg, A.F., Oehlers, S.H., Saelens, J.W., Sisk, D.M., Jurcic Smith, K.L., Lee, S., Tobin, D.M., 2015. CLARITY and PACT-based imaging of adult zebrafish and mouse for whole-animal analysis of infections. Dis Model Mech 8, 1643–1650.

Davis, J.M., Clay, H., Lewis, J.L., Ghori, N., Herbomel, P., Ramakrishnan, L., 2002. Real-time visualization of mycobacterium-macrophage interactions leading to initiation of granuloma formation in zebrafish embryos. Immunity 17, 693–702.

Davis, J.M., Ramakrishnan, L., 2009. The role of the granuloma in expansion and dissemination of early tuberculous infection. Cell 136, 37–49.

Ellett, F., Lieschke, G.J., 2012. Computational quantification of fluorescent leukocyte numbers in zebrafish embryos. Methods Enzymol 506, 425–435.

Eruslanov, E.B., Lyadova, I.V., Kondratieva, T.K., Majorov, K.B., Scheglov, I.V., Orlova, M.O., Apt, A.S., 2005. Neutrophil responses to Mycobacterium tuberculosis infection in genetically susceptible and resistant mice. Infect Immun 73, 1744–1753.

Eum, S.Y., Kong, J.H., Hong, M.S., Lee, Y.J., Kim, J.H., Hwang, S.H., Cho, S.N., Via, L.E., Barry, C.E., 3rd, 2010. Neutrophils are the predominant infected phagocytic cells in the airways of patients with active pulmonary TB. Chest 137, 122–128.

Hall, C., Flores, M.V., Storm, T., Crosier, K., Crosier, P., 2007. The zebrafish lysozyme C promoter drives myeloid-specific expression in transgenic fish. BMC Dev Biol 7, 42.

Hui, S.P., Sheng, D.Z., Sugimoto, K., Gonzalez-Rajal, A., Nakagawa, S., Hesselson, D., Kikuchi, K., 2017. Zebrafish Regulatory T Cells Mediate Organ-Specific Regenerative Programs. Dev Cell 43, 659–672 e655.

Jin, S.W., Beis, D., Mitchell, T., Chen, J.N., Stainier, D.Y., 2005. Cellular and molecular analyses of vascular tube and lumen formation in zebrafish. Development 132, 5199–5209.

Johansen, M.D., Kasparian, J.A., Hortle, E., Britton, W.J., Purdie, A.C., Oehlers, S.H., 2018. Mycobacterium marinum infection drives foam cell differentiation in zebrafish infection models. Dev Comp Immunol 88, 169–172.

Lin, H.F., Traver, D., Zhu, H., Dooley, K., Paw, B.H., Zon, L.I., Handin, R.I., 2005. Analysis of thrombocyte development in CD41-GFP transgenic zebrafish. Blood 106, 3803–3810.

Marjoram, L., Alvers, A., Deerhake, M.E., Bagwell, J., Mankiewicz, J., Cocchiaro, J.L., Beerman, R.W., Willer, J., Sumigray, K.D., Katsanis, N., Tobin, D.M., Rawls, J.F., Goll, M.G., Bagnat, M., 2015. Epigenetic control of intestinal barrier function and inflammation in zebrafish. Proc Natl Acad Sci U S A 112, 2770–2775.

Oehlers, S.H., Cronan, M.R., Scott, N.R., Thomas, M.I., Okuda, K.S., Walton, E.M., Beerman, R.W., Crosier, P.S., Tobin, D.M., 2015. Interception of host angiogenic signalling limits mycobacterial growth. Nature 517, 612–615.

Parikka, M., Hammaren, M.M., Harjula, S.K., Halfpenny, N.J., Oksanen, K.E., Lahtinen, M.J., Pajula, E.T., Iivanainen, A., Pesu, M., Ramet, M., 2012. Mycobacterium marinum Causes a Latent Infection that Can Be Reactivated by Gamma Irradiation in Adult Zebrafish. PLoS Pathog 8, e1002944.

Prouty, M.G., Correa, N.E., Barker, L.P., Jagadeeswaran, P., Klose, K.E., 2003. Zebrafish-Mycobacterium marinum model for mycobacterial pathogenesis. FEMS Microbiol. Lett. 225, 177–182.

Ramakrishnan, L., 2012. Revisiting the role of the granuloma in tuberculosis. Nat Rev Immunol 12, 352–366.

Risalde, M.A., Lopez, V., Contreras, M., Mateos-Hernandez, L., Gortazar, C., de la Fuente, J., 2018. Control of mycobacteriosis in zebrafish (Danio rerio) mucosally vaccinated with heat-inactivated Mycobacterium bovis. Vaccine 36, 4447–4453.

Schindelin, J., Arganda-Carreras, I., Frise, E., Kaynig, V., Longair, M., Pietzsch, T., Preibisch, S., Rueden, C., Saalfeld, S., Schmid, B., Tinevez, J.Y., White, D.J., Hartenstein, V., Eliceiri, K., Tomancak, P., Cardona, A., 2012. Fiji: an open-source platform for biological-image analysis. Nat Methods 9, 676–682.

Schneider, C.A., Rasband, W.S., Eliceiri, K.W., 2012. NIH Image to ImageJ: 25 years of image analysis. Nat Methods 9, 671–675.

Sugimoto, K., Hui, S.P., Sheng, D.Z., Nakayama, M., Kikuchi, K., 2017. Zebrafish FOXP3 is required for the maintenance of immune tolerance. Dev Comp Immunol 73, 156–162.

Swaim, L.E., Connolly, L.E., Volkman, H.E., Humbert, O., Born, D.E., Ramakrishnan, L., 2006. Mycobacterium marinum infection of adult zebrafish causes caseating granulomatous tuberculosis and is moderated by adaptive immunity. Infect Immun 74, 6108–6117.

Takaki, K., Cosma, C.L., Troll, M.A., Ramakrishnan, L., 2012. An in vivo platform for rapid high-throughput antitubercular drug discovery. Cell reports 2, 175–184.

Takaki, K., Davis, J.M., Winglee, K., Ramakrishnan, L., 2013. Evaluation of the pathogenesis and treatment of Mycobacterium marinum infection in zebrafish. Nat Protoc 8, 1114–1124.

Tenor, J.L., Oehlers, S.H., Yang, J.L., Tobin, D.M., Perfect, J.R., 2015. Live Imaging of Host-Parasite Interactions in a Zebrafish Infection Model Reveals Cryptococcal Determinants of Virulence and Central Nervous System Invasion. mBio 6.

van der Sar, A.M., Abdallah, A.M., Sparrius, M., Reinders, E., Vandenbroucke-Grauls, C.M., Bitter, W., 2004. Mycobacterium marinum strains can be divided into two distinct types based on genetic diversity and virulence. Infect Immun 72, 6306–6312.

van Leeuwen, L.M., van der Kuip, M., Youssef, S.A., de Bruin, A., Bitter, W., van Furth, A.M., van der Sar, A.M., 2014. Modeling tuberculous meningitis in zebrafish using Mycobacterium marinum. Dis Model Mech 7, 1111–1122.

Volkman, H.E., Clay, H., Beery, D., Chang, J.C., Sherman, D.R., Ramakrishnan, L., 2004. Tuberculous granuloma formation is enhanced by a mycobacterium virulence determinant. PLoS Biol 2, e367.

Weerdenburg, E.M., Abdallah, A.M., Mitra, S., de Punder, K., van der Wel, N.N., Bird, S., Appelmelk, B.J., Bitter, W., van der Sar, A.M., 2012. ESX-5-deficient Mycobacterium marinum is hypervirulent in adult zebrafish. Cell Microbiol 14, 728–739.

