## Supplementary material for "High content analysis of granuloma histology and neutrophilic inflammation in adult zebrafish infected with *Mycobacterium marinum*": Macro template

The imagej macro below will remove scale information, set a threshold limit of 20-255, and then count up to 70 regions of interest presenting a pixel count of fluorescence reporter area in each granuloma.

run("8-bit");

run("Set Scale...", "distance=0");

setAutoThreshold("Default dark");

setThreshold(20, 255);

roiCount = roiManager("count");

for (i=0; i<roiCount; i++) {

roiManager("select", i);

run("Analyze Particles...", "summarize");

roiManager("select", i+1);

run("Analyze Particles...", "summarize");

roiManager("select", i+2);

run("Analyze Particles...", "summarize");

roiManager("select", i+3);

run("Analyze Particles...", "summarize");

roiManager("select", i+4);

run("Analyze Particles...", "summarize");

roiManager("select", i+5);

run("Analyze Particles...", "summarize");

roiManager("select", i+6);

run("Analyze Particles...", "summarize");

roiManager("select", i+7);

run("Analyze Particles...", "summarize");

roiManager("select", i+8);

run("Analyze Particles...", "summarize");

roiManager("select", i+9);

run("Analyze Particles...", "summarize");

roiManager("select", i+10);

run("Analyze Particles...", "summarize");

roiManager("select", i+11);

run("Analyze Particles...", "summarize");

roiManager("select", i+12);

run("Analyze Particles...", "summarize");

roiManager("select", i+13);

run("Analyze Particles...", "summarize");

roiManager("select", i+14);

run("Analyze Particles...", "summarize");

roiManager("select", i+15);

run("Analyze Particles...", "summarize");

roiManager("select", i+16);

run("Analyze Particles...", "summarize");

roiManager("select", i+17);

run("Analyze Particles...", "summarize");

roiManager("select", i+18);

run("Analyze Particles...", "summarize");

roiManager("select", i+19);

run("Analyze Particles...", "summarize");

roiManager("select", i+20);

run("Analyze Particles...", "summarize");

roiManager("select", i+21);

run("Analyze Particles...", "summarize");

roiManager("select", i+22);

run("Analyze Particles...", "summarize");

roiManager("select", i+23);

run("Analyze Particles...", "summarize");

roiManager("select", i+24);

run("Analyze Particles...", "summarize");

roiManager("select", i+25);

run("Analyze Particles...", "summarize");

roiManager("select", i+26);

run("Analyze Particles...", "summarize");

roiManager("select", i+27);

run("Analyze Particles...", "summarize");

roiManager("select", i+28);

run("Analyze Particles...", "summarize");

roiManager("select", i+29);

run("Analyze Particles...", "summarize");

roiManager("select", i+30);

run("Analyze Particles...", "summarize");

roiManager("select", i+31);

run("Analyze Particles...", "summarize");

roiManager("select", i+32);

run("Analyze Particles...", "summarize");

roiManager("select", i+33);

run("Analyze Particles...", "summarize");

roiManager("select", i+34);

run("Analyze Particles...", "summarize");

roiManager("select", i+35);

run("Analyze Particles...", "summarize");

roiManager("select", i+36);

run("Analyze Particles...", "summarize");

roiManager("select", i+37);

run("Analyze Particles...", "summarize");

roiManager("select", i+38);

run("Analyze Particles...", "summarize");

roiManager("select", i+39);

run("Analyze Particles...", "summarize");

roiManager("select", i+40);

run("Analyze Particles...", "summarize");

roiManager("select", i+41);

run("Analyze Particles...", "summarize");

roiManager("select", i+42);

run("Analyze Particles...", "summarize");

roiManager("select", i+43);

run("Analyze Particles...", "summarize");

roiManager("select", i+44);

run("Analyze Particles...", "summarize");

roiManager("select", i+45);

run("Analyze Particles...", "summarize");

roiManager("select", i+46);

run("Analyze Particles...", "summarize");

roiManager("select", i+47);

run("Analyze Particles...", "summarize");

roiManager("select", i+48);

run("Analyze Particles...", "summarize");

roiManager("select", i+49);

run("Analyze Particles...", "summarize");

roiManager("select", i+50);

run("Analyze Particles...", "summarize");

roiManager("select", i+51);

run("Analyze Particles...", "summarize");

roiManager("select", i+52);

run("Analyze Particles...", "summarize");

roiManager("select", i+53);

run("Analyze Particles...", "summarize");

roiManager("select", i+54);

run("Analyze Particles...", "summarize");

roiManager("select", i+55);

run("Analyze Particles...", "summarize");

roiManager("select", i+56);

run("Analyze Particles...", "summarize");

roiManager("select", i+57);

run("Analyze Particles...", "summarize");

roiManager("select", i+58);

run("Analyze Particles...", "summarize");

roiManager("select", i+59);

run("Analyze Particles...", "summarize");

roiManager("select", i+60);

run("Analyze Particles...", "summarize");

roiManager("select", i+61);

run("Analyze Particles...", "summarize");

roiManager("select", i+62);

run("Analyze Particles...", "summarize");

roiManager("select", i+63);

run("Analyze Particles...", "summarize");

roiManager("select", i+64);

run("Analyze Particles...", "summarize");

roiManager("select", i+65);

run("Analyze Particles...", "summarize");

roiManager("select", i+66);

run("Analyze Particles...", "summarize");

roiManager("select", i+67);

run("Analyze Particles...", "summarize");

roiManager("select", i+68);

run("Analyze Particles...", "summarize");

roiManager("select", i+69);

run("Analyze Particles...", "summarize");

roiManager("select", i+70);
